# Rising SARS-CoV-2 Seroprevalence and Patterns of Cross-Variant Antibody Neutralization in UK Domestic Cats

**DOI:** 10.1101/2022.11.18.517046

**Authors:** Grace B Tyson, Sarah Jones, Nicola Logan, Michael McDonald, Pablo R Murcia, Brian J Willett, William Weir, Margaret J Hosie

## Abstract

Recent evidence confirming cat-to-human SARS-CoV-2 transmission has highlighted the importance of monitoring infection in domestic cats. Although the effects of SARS-CoV-2 infection on feline health are poorly characterized, cats have close contact with humans, and with both domesticated and wild animals. Accordingly, they could act as a reservoir of infection, an intermediate host and a source of novel variants. To investigate the spread of the virus in the cat population, serum samples were tested for SARS-CoV-2 antibodies by ELISA and a pseudotype-based virus neutralization assay, designed to detect exposure to variants known to be circulating in the human population. Overall seroprevalence was 3.2%, peaking at 5.3% in autumn 2021. Variant-specific neutralizing antibody responses were detected with titers waning over time. The variant-specific response in the feline population correlated with and trailed the variants circulating in the human population, indicating multiple ongoing human-to-cat spill-over events.

## Introduction

Throughout the COVID-19 pandemic, there have been sporadic cases of SARS-CoV-2 infection detected in felids, particularly in domestic cats (*1, 2, 3, 4, 5*). SARS-CoV-2 infections in domestic cats have been reported in the UK (*6, 7*) and over 20 other countries worldwide, with global spread likely to be considerably greater. Infections have also been reported in several other felids including snow leopards, lions, tigers and fishing cats (*4, 8*). The ACE2 receptor molecule that facilitates SARS-CoV-2 cell entry is well conserved across many mammalian species (*9, 10*). The amino acid sequence of the ACE2 protein of *Felis catus*, is highly similar to human ACE2 and this may contribute to the high susceptibility of felids to SARS-CoV-2 infection (*3*).

Despite current evidence showing most cases of SARS-CoV-2 in felids are spillover infections resulting from close contact with infected humans (*11*), SARS-CoV-2-specific antibodies have been found in stray cats in Rio De Janeiro (*12*), and in abandoned cats in Wuhan, indicating cats might be infected from other sources (*13*). Similarly, cat-to-cat transmission has been demonstrated experimentally (*14, 15, 16*).

Recently, a case of cat-to-human SARS-CoV-2 transmission was observed in Thailand, which was indicated by comparing viral genome sequences from the cat and its attending veterinary surgeon (*17*). Given that domestic cats are frequently in close contact with humans, if they become a reservoir for the virus, they could initiate new outbreaks or re-introduce SARS-CoV-2 into humans (*18*). Moreover, if SARS-CoV-2 adapts to replicate more efficiently in cats, they could contribute to the emergence of novel variants. It has been suggested that the Omicron variant might have emerged from a cross-species transmission of SARS-CoV-2 into an animal reservoir, in which mutations accumulated, then spilled back to humans (*19*). This pattern of variant emergence was observed during the 2020 outbreak of SARS-CoV-2 on Dutch mink farms (*20*). Infection of stray cats living on a mink farm, suggestive of mink-to-cat transmission, has also been reported (*21, 22*).

Several clinical outcomes of feline SARS-CoV-2 infection have been documented (*23*), from asymptomatic infections (*24*) to mild respiratory signs (*25*). More severe sequelae include myocarditis (*26, 27*), which can be severe and lead to death or necessitate euthanasia. Estimating the frequency of asymptomatic infections by RT-qPCR is technically challenging as there is a narrow window of positivity (*28*). Cui *et al* (2020) suggested cats might be less likely to display signs than humans as two key sensory components of the inflammasome pathway, Aim2 and NLRP1, are absent in both domestic cats and tigers (*29*). It was hypothesized that this might confer an evolutionary advantage of a reduction in excessive cytokine release, resulting in less host tissue damage and milder inflammatory symptoms during SARS-CoV-2 infection in these animals.

Despite the potential impact of SARS-CoV-2 on feline health, there is currently no official surveillance program for monitoring SARS-CoV-2 infection or exposure in UK cats. Diagnostic RT-qPCR testing has primarily been undertaken by researchers and has been constrained by a narrow case definition by the UK’s competent authority(*30*).

It has been demonstrated experimentally that domestic cats mount a neutralizing antibody response against SARS-CoV-2 that prevents re-infection from a second viral challenge (*16*) and a feline IgG response has been detected against both the nucleocapsid and spike proteins via ELISA (*31, 32*). Cats have also been found to produce a neutralizing antibody response against multiple SARS-CoV-2 variants (*33*).

The antibody response to both SARS-CoV-2 infection and vaccination in humans wanes over time, more rapidly than for other human coronavirus infections, allowing for re-infection with SARS-CoV-2 (*34, 35*). It has also been found that less severe clinical outcomes (*36*) and longer-lived immunity is exhibited by children than adults in response to SARS-CoV-2 infection (*37*). However, it is unknown if feline SARS-CoV-2 immunity is transient or if age-dependent immune longevity and clinical outcomes are also a feature of feline infections.

In humans, virus neutralizing antibodies generated in response to SARS-CoV-2 vaccines, currently based on the ancestral Wuhan-Hu-1 strain, are less effective against the Delta and Omicron variants (*38, 39, 40, 41, 42*). A cat that has been infected with one variant might resist re-infection with the same variant but remain susceptible to infection with a different variant, similar to the phenomenon observed in humans (*43*).

There are many breeds of domesticated cat, and it is possible genetic differences generated by selective breeding could have an impact on immunity (*44*), susceptibility to infection or the severity of clinical signs, whether by selection for a genetic defect or narrowing of major histocompatibility complex (MHC) diversity. For example, pedigree cats are more likely to develop feline infectious peritonitis following feline coronavirus infection than non-pedigree cats (*45*).

However, it should be appreciated that the breeding of pedigree cats is often associated with other risk factors such as multi-cat households and being kept indoors, and the actual genetic basis for susceptibility has not been quantified.

The aim of the present study was to assess the seroprevalence of SARS-CoV-2 infection in UK cats during the COVID-19 pandemic, using an ELISA to measure antibodies recognizing the receptor binding domain of the SARS-CoV-2 S-protein and a pseudotype-based neutralization assay to measure titers of virus neutralizing antibodies. Neutralizing titers were measured against a panel of HIV (SARS-CoV-2) pseudotypes bearing the S protein of the predominant circulating variants in the UK to investigate the specificity of the neutralizing response and whether it correlated with the variants likely to have been circulating at the time of infection.

## Methods

### Samples

Residual blood samples for serological testing were obtained from the University of Glasgow Veterinary Diagnostic Services laboratory (VDS). These samples had been submitted by practicing UK veterinary surgeons for purposes including routine monitoring, pre-breeding screening, testing for other infections and the diagnosis of hormonal disorders (Fig 8). Residual serum/plasma that would otherwise have been discarded after all requested tests had been completed was used for this study. None of the samples had been submitted because of suspected SARS-CoV-2 infection. These samples represented a cohort broadly representative of the domestic cat population throughout the UK. Poor quality samples, for example those displaying marked hemolysis, were excluded. Ethical approval for the study and was granted by the University of Glasgow Veterinary Ethics Committee (EA27/20). Samples were given a unique identification number on arrival, and investigators (GT, NL and SJ) were blinded to sample metadata until the data analysis stage.

### Serological testing

Samples were initially screened at a final dilution of 1 in 100 using a pseudotype-based virus neutralization assay (PVNA). PVNA positive samples were confirmed using a double antigen binding assay (DABA) ELISA that detected antibodies recognizing the receptor-binding domain of the SARS-CoV-2 S protein. Neutralizing antibody titers were estimated by performing a PVNA with serially diluted samples.

For the neutralization assays, HIV (SARS-CoV-2) pseudotypes were constructed bearing the spike proteins of either the Wuhan-Hu-1 D614G (B.1), Alpha (B.1.1.7), Delta (B.1.617.2) or Omicron (BA.1) SARS-CoV-2 variants. Samples collected early in the pandemic were tested against Wuhan-Hu-1 D614G (B.1) only while new variants were included in the assay over time, as each new SARS-CoV-2 variant emerged during subsequent waves of the pandemic.

### Pseudotype-based Virus Neutralization Assay

The method for this assay has been described previously (*40*). Briefly, HEK293, HEK293T, and 293-ACE2 cells were maintained in Dulbecco’s modified Eagle’s medium (DMEM) supplemented with 10% fetal bovine serum, 200mM L-glutamine, 100μg/ml streptomycin and 100 IU/ml penicillin. HEK293T cells were transfected with the appropriate SARS-CoV-2 S gene expression vector (wild type or variant) in conjunction with p8.91 (*46*) and pCSFLW (*47*) using polyethylenimine (PEI, Polysciences, Warrington, USA). HIV (SARS-CoV-2) pseudotypes were harvested from culture fluids 48 hours post-transfection, filtered at 0.45μm, aliquoted and frozen at −80°C prior to use. The SARS-CoV-2 spike glycoprotein expression constructs were synthesized by GenScript (Netherlands). Constructs bore the following mutations relative to the Wuhan-Hu-1 sequence (GenBank: MN908947):

- **B.1 (Wuhan-Hu-1 D614G**) – D614G
- **B.1.1.7 (Alpha)** – Δ69-70, Δ144, N501Y, A570D, D614G, P681H, T716I, S982A, D1118H
- **B.1.617.2 (Delta)** – T19R, G142D, Δ156-157, R158G, L452R, T478K, D614G, P681R, D950N
- **B.1.1.529 (Omicron BA.1)** - A67V, Δ69-70, T95I, G142D/Δ143-145, Δ211/L212I, ins214EPE, G339D, S371L, S373P, S375F, K417N, N440K, G446S, S477N, T478K, E484A, Q493R, G496S, Q498R, N501Y, Y505H, T547K, D614G, H655Y, N679K, P681H, N764K, D796Y, N856K, Q954H, N969K, L981F

All synthesized S genes were codon-optimized, incorporated the mutation K1255STOP to enhance surface expression, and were cloned into the pcDNA3.1(+) eukaryotic expression vector. 293-ACE2 target cells (*48*) were maintained in complete DMEM supplemented with 2μg/ml puromycin.

The fixed dilution screen was performed with serum/plasma diluted 1:50 in complete DMEM (in duplicate) for each pseudotype. Diluted samples were incubated with HIV (SARS-CoV-2) pseudotypes for 1 hour and plated onto 239-ACE2 target cells. After 48-72 hours, luciferase activity was quantified by the addition of Steadylite Plus chemiluminescence substrate and analysis on a Perkin Elmer EnSight multimode plate reader (PerkinElmer, Beaconsfield, UK). Samples which reduced the infectivity of the pseudotypes by at least 90% were classed as positive. For positive samples, neutralizing activity was then quantified by serial dilution. Each sample was serially diluted (in triplicate) from 1:50 to 1:36450 in complete DMEM prior to incubation with the respective viral pseudotype. Antibody titer was then estimated by interpolating the point at which infectivity had been reduced to 90% of the value for the no serum control samples.

Seropositive cats were categorized according to the pseudotype variant against which the highest neutralizing titer was obtained. For example, samples showing a higher titer against the Delta pseudotype compared to the other pseudotypes were categorized as “Delta dominant”.

### Double Antigen Bridging Assay ELISA

All samples that appeared positive on the initial fixed dilution PVNA were tested using a species agnostic double antigen bridging assay (Microimmune SARS-CoV-2 Double Antigen Bridging Assay (COVT016), Clin-Tech, Guildford, England) according to the manufacturer’s instructions, to determine whether samples contained antibodies to the B.1 SARS-CoV-2 receptor-binding domain. This was used to confirm results of the pseudotype-based neutralization assay by confirming low chemiluminescence readings were caused by high levels of antibody rather than any toxic contamination of samples killing the cells.

### Data Analysis

Duplicate samples were removed while samples from the same animal tested multiple times were identified and the earliest sample was used to estimate seroprevalence. A small number of animals had multiple samples submitted to the VDS at different times and, using these samples, longitudinal titers were tabulated to explore the effect of time on the development of the humoral response to SARS-CoV-2. Data were analyzed and graphs prepared using GraphPad Prism 9.3.1. and Microsoft Excel. Distribution of data was assessed using a Shapiro-Wilk Normality test. Sample metadata (age, sex, breed, location) was acquired from information recorded in the VDS database, which was supplied by submitting veterinary surgeons Differences between groups were assessed for significance in paired data using a Wilcoxon test and in unpaired data using a Mann-Whitney test. Significance of categorical data was assessed using a Chi-Square test.

## Results

### Sample population

Serum samples from 2309 different cats sampled between the 21^st^ of April 2020 and the 7^th^ of February 2022 were tested (Fig. 1a). Within this sample group, 1174 (50.9%) were male, 853 (36.9%) were female, while the sex of the cat was not recorded for the remaining 282 (12.2%). The ages of the cats ranged from <1 to 21 years (mean 5.1 years, median 3 years). Age data were not included on the submission forms for 350 (15.1%) of animals tested. The group comprised 56% non-pedigree cats (1300/2309) and 31% pedigree cats (720/2309), with the remainder, 13% (289/2309), being of unstated breed.

**Figure 1:**
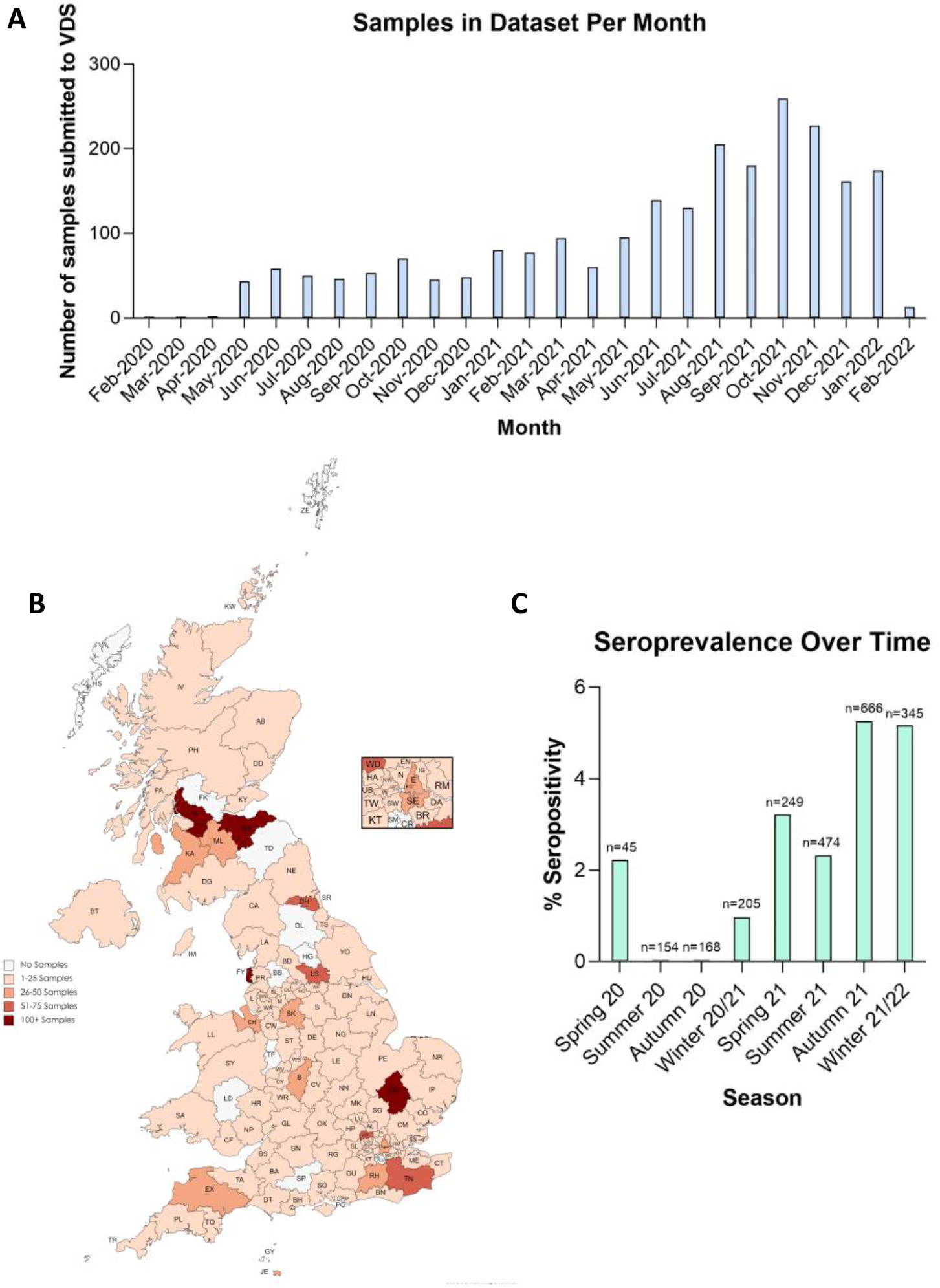
Overview of Samples included in Analysis. The number of samples tested per month (A). The location of the veterinary practices that submitted samples used in this study (B). The percentage seropositivity of samples per 3-month period and sample size for each period. (C). Overall seropositivity across all samples was 3.2% (75/2309).

The study included samples from 112 of the UK’s 126 postcode areas, with an uneven distribution amongst postcode areas that was unrelated to the local human population density. Overrepresented areas included Blackpool, Glasgow, Edinburgh and Cambridge (Fig 1b).

### Overall seroprevalence

Seroprevalence of SARS-CoV-2 antibodies in UK cats increased over time (Fig 1c). Overall, the seroprevalence during the study period was 3.2% (75/2309, 95% CI = 2.56%-4.05%). The seroprevalence was highest during the periods Sep-Nov 2021 (35/666, 5.3%, 95% CI = 3.69%-7.23%) and Dec 2021-Feb 2022 (18/348, 5.2%, 95% CI = 3.09%-8.05%).

### Seroprevalence amongst different groups

A greater proportion of pedigree cats (31/720, 4.3%, 95% CI = 2.94%-6.06%) than non-pedigree cats (39/1300, 3%, 95% CI = 2.14%-4.08%) tested seropositive for SARS-CoV-2 (Fig 2a & 2b), with Bengal, Siamese and British Blue/Shorthair breeds showing the highest seroprevalence, however, the differences in seroprevalence between different breeds were not significant (p=0.07) (Fig 2c). Maine Coon cats were the only breed with over 30 cats sampled that showed a lower seroprevalence than the population mean. The strength of VNA titers elicited by pedigree and non-pedigree cats were not found to be significantly different (p=0.5) (Fig 2d). A greater proportion of male than female cats in this study were seropositive, however, this was not significant (p=0.5) (Fig 3) and there was no significant difference between the average highest titer for each sex group (p=0.7). There was also no significant difference in cat age between positive and negative samples (p=0.89) (Fig 4), nor any correlation between age and neutralization titer (Fig 4).

**Figure 2:**
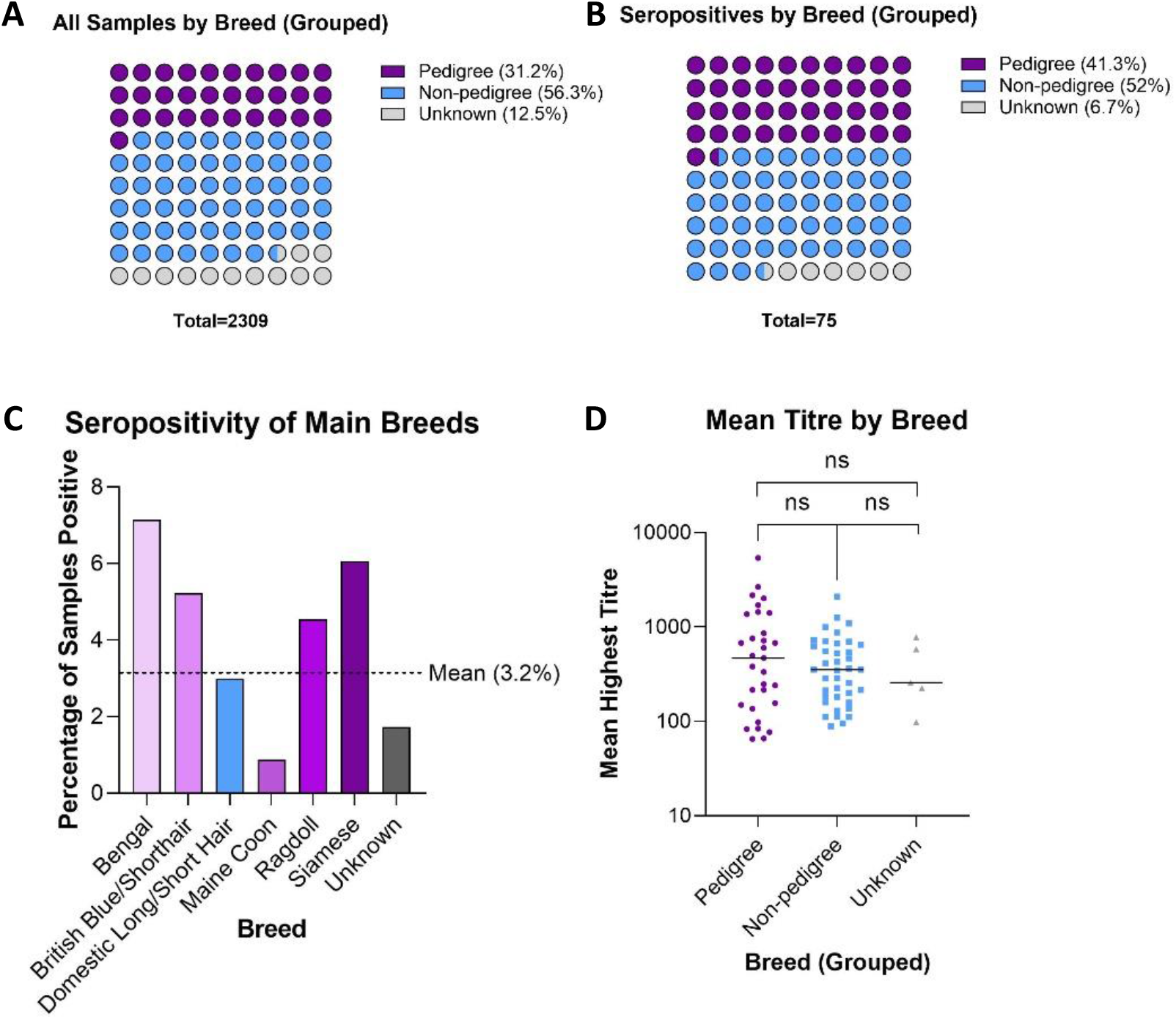
Total and seropositive samples analysed by breed. Total samples in this study classified by breed; Pedigree, non-pedigree or unknown (A). Seropositive samples classified by breed (B). The percentage seropositivity for each breed with over 30 samples included in the study (C). The highest virus neutralization titer of each seropositive sample categorized by breed (D). There was no significant difference between the highest titres of each breed category.

**Figure 3:**
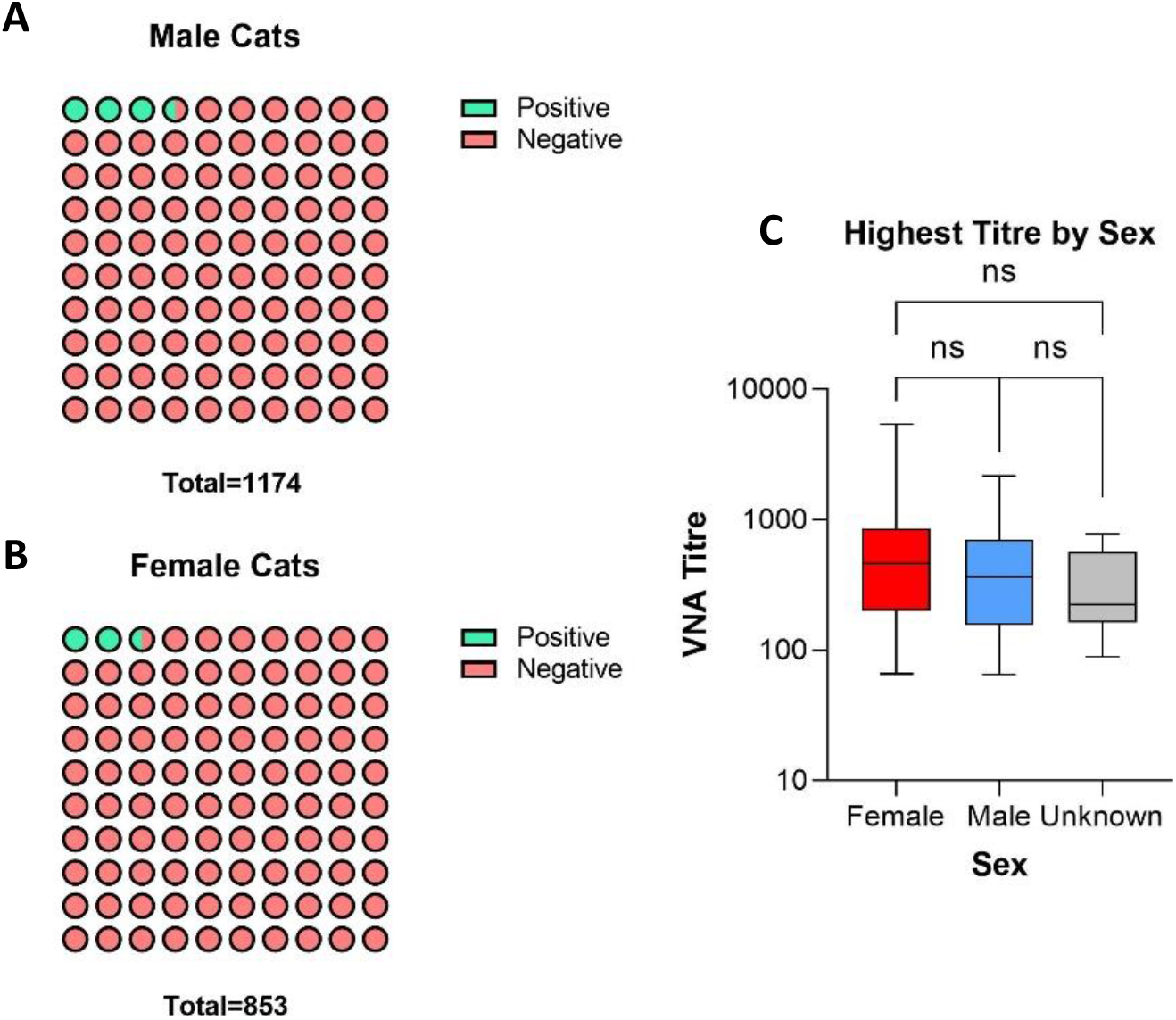
Overview of sex of seropositive animals. Total samples tested categorized by sex (A,B). The highest titer of animals in each sex category was not significantly different (Mann-Whitney test) (C).

**Figure 4:**
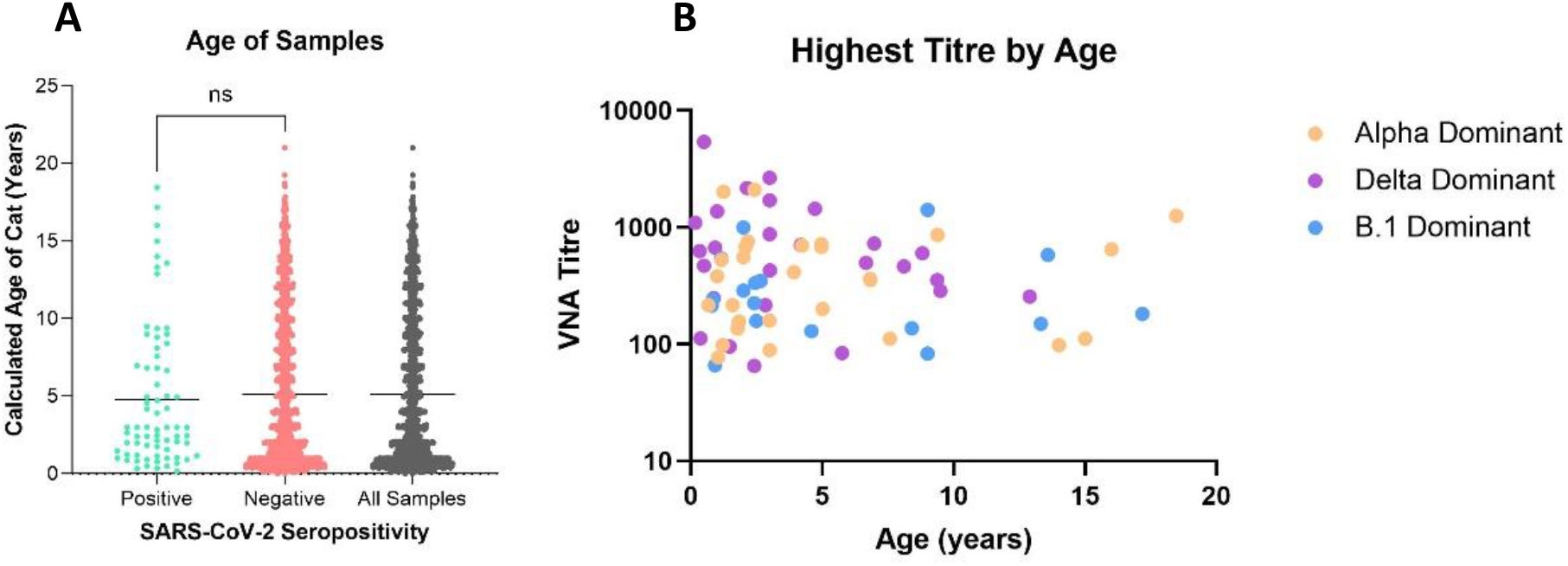
Overview of age of seropositive animals. The age distribution of positive and negative samples analyzed using a Mann-Whitney test (A). No significant difference was seen between the average age of positive and negative samples. Each sample’s highest titre plotted against the age of the cat sampled (B). No correlation was observed.

### Antibody titers against pseudotypes of different SARS-CoV-2 variants

A comparison of the specificity of the neutralizing response suggested 27/75 (36%) of seropositive cats in this study displayed responses that were “Delta dominant”, 31/75 (41.3%) were “Alpha dominant” and 17/75 (22.7%) were “B.1 dominant”. On average, Delta dominant cats displayed higher neutralization titers (mean: 760) against their dominant pseudotype compared to Alpha (488, p=0.06) or B.1 (329, p=0.02) dominant cats (Fig 5). Throughout the time-period of sampling in this study (April 2020-Feb 2022), no Omicron dominant seropositive cats were identified.

**Figure 5:**
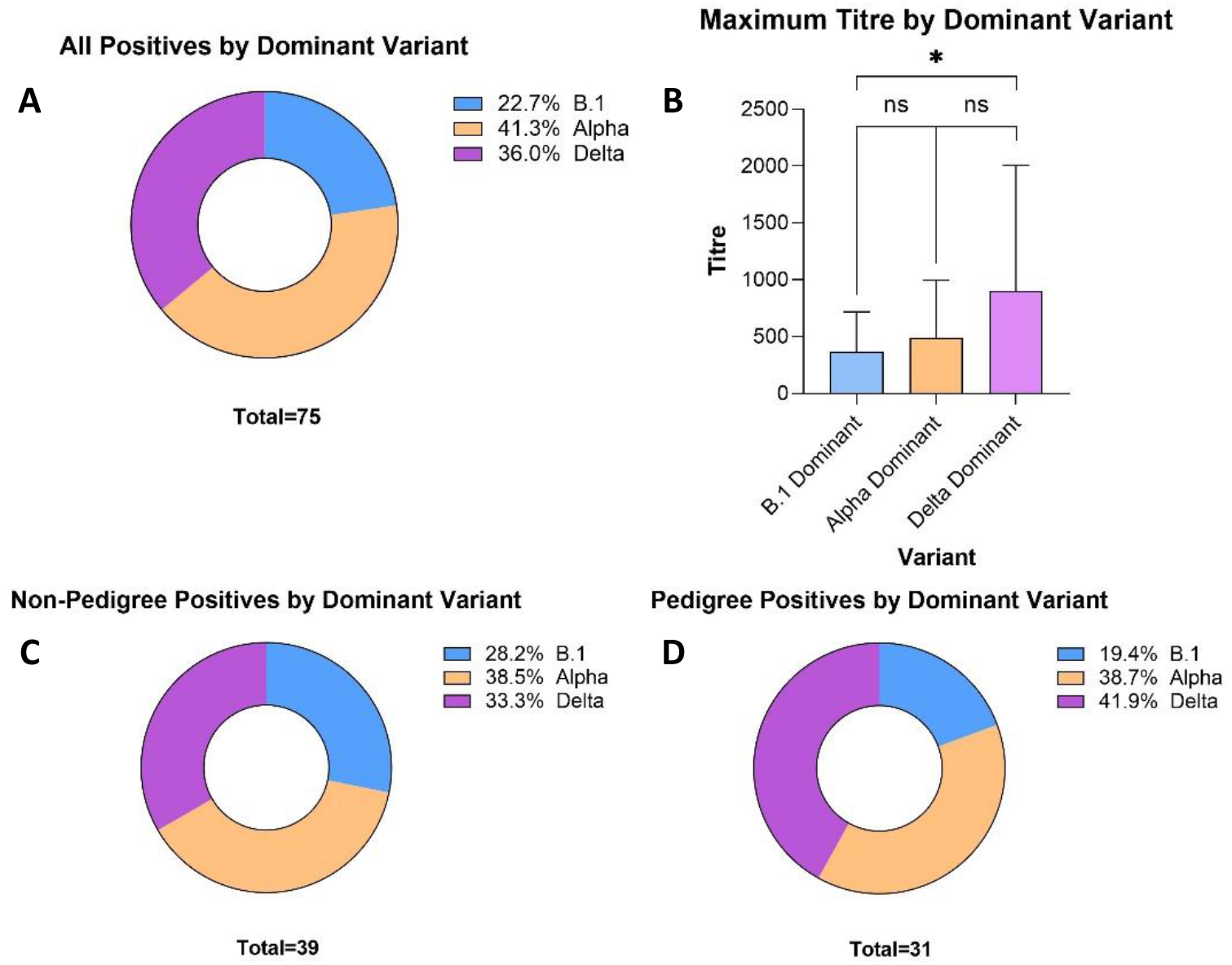
Seropositive cases shown by dominant variant. Seropositive samples categorized by their dominant variant (A). The average titre produced by each serum sample against its dominant variant (B). Normality of sample distribution was assessed using a Shapiro-wilk test and significance was assessed using a Mann-Whitney test (ns= not significant, *= p<0.05). Seropositive samples categorized by breed – either non-pedigree (C) or pedigree (D). Seropositive cats of unknown breed were not included in this figure.

A greater proportion of pedigree than non-pedigree cats were found to be Delta dominant but this was not significant (p=0.4). Non-pedigree cats showed a more even distribution of cases by variant, whereas comparatively few pedigree cats were infected with the B.1 variant (Fig 5).

There appears to be a correlation between the dominant variant observed in cats and the timeline of variant emergence into the human population. Detection of new dominant variants in cats trails the detection of the variant in the humans, however, dominant titers were still detected against extinct variants long after human cases had subsided (Fig. 6).

**Figure 6:**
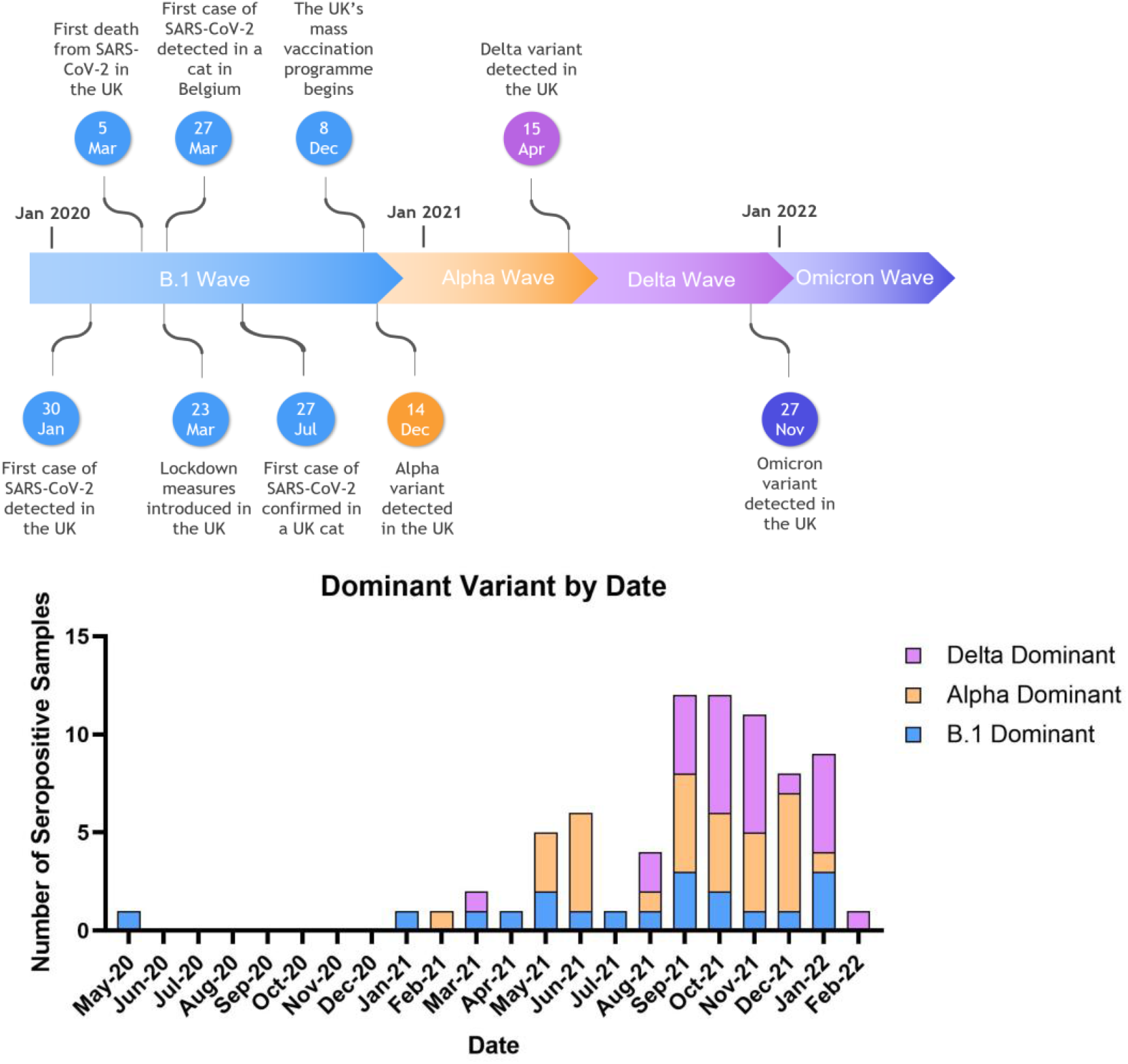
Overview of dominant variant of seropositive samples by date. Seropositive samples categorized by dominant variant and plotted by month. Results are displayed as a percentage of all seropositive samples from that period. Also shown is a timeline of key events of the COVID-19 pandemic in the UK and emergence of major variants into the UK human population.

Distinct patterns of neutralization were observed in that B.1 dominant samples generally had slightly lower titers against the Alpha pseudotype than against B.1, but significantly lower titers against the Delta pseudotype, and significantly lower still against Omicron. Alpha dominant samples showed slightly lower B.1 titers than Alpha titers and markedly lower Delta and Omicron titers. Delta dominant cats showed similar titers against the B.1, Alpha and Omicron pseudotypes, all of which were markedly lower than their Delta titers (Fig 7).

**Figure 7:**
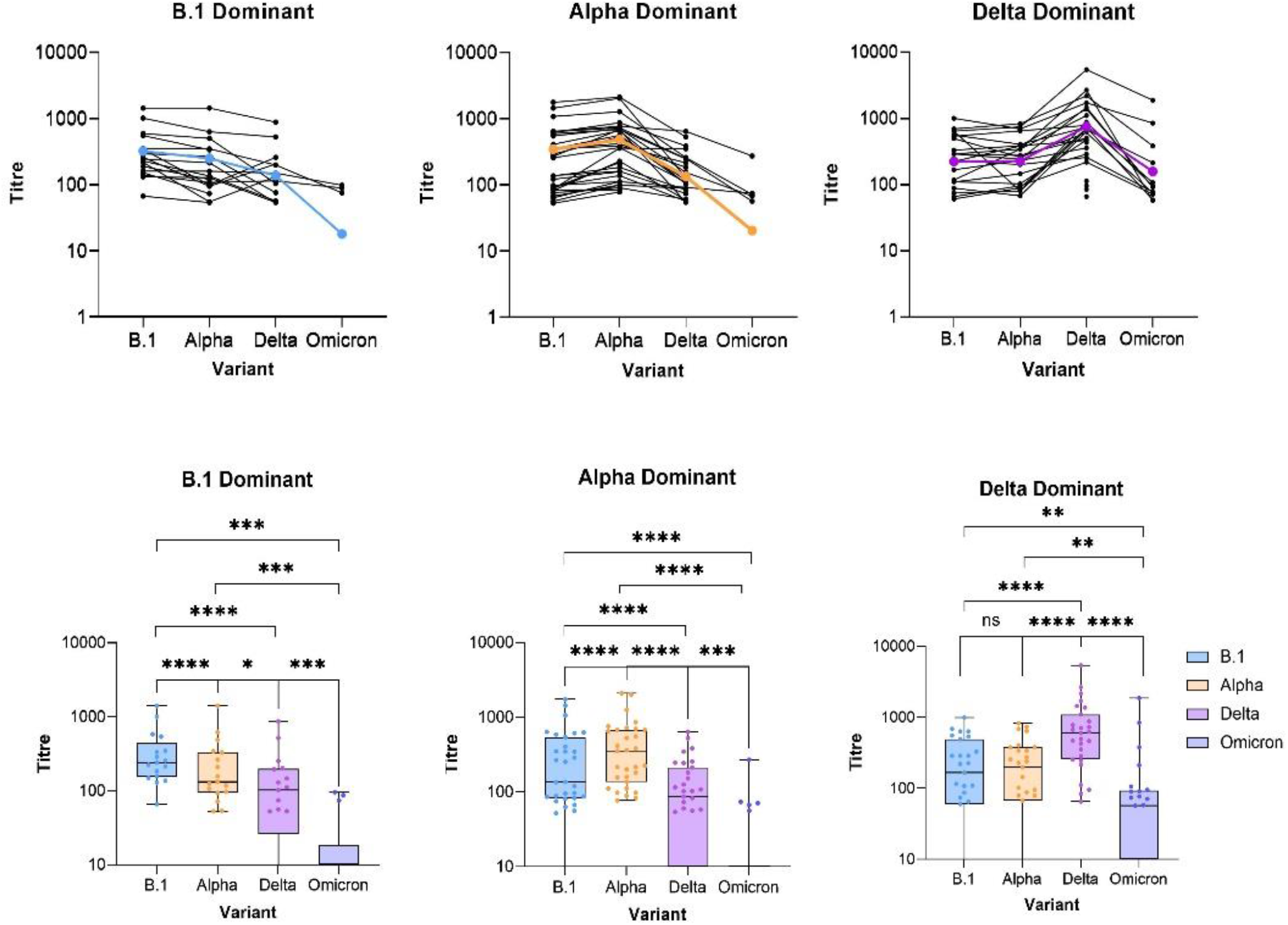
Virus neutralisation titres of seropositive samples grouped by dominant variant. Neutralizing titers for samples classified by dominant variant, showing the three distinct patterns of immunity (ns= not significantly different, asterisks indicate significant differences as follows: *= p<0.05, ** = p<0.01, *** = p<0.001, ****= p<0.0001, Wilcoxon test). Mean patterns of cross-neutralization for each dominant group are displayed in bold color in line graphs

### Longitudinal samples

Five seropositive cats had samples taken at least 12 days apart. In all five cases, it was observed that neutralizing titers against SARS-CoV-2 waned over time. Percentage decrease in titer per day was highly variable across samples, but in the case of three of the cats, was consistent across all variants (Table 1).

**Table 1:** Overview of longitudinal samples. Five animals had two samples submitted to the study taken ≥12 days apart. The earliest sample was used in the overall analysis, however, newer samples were also tested and the titres against each variant are shown for each sample with the earlier sample on top and later below. Titres are colour-coded by size. Percentage change in titre per day is also shown.

| Sample | Days<br>Between<br>Sampling | Titres |  |  | Percentage change per day |  |  |
| --- | --- | --- | --- | --- | --- | --- | --- |
|  |  | *B.1* | *Alpha* | *Delta* | *B.1* | *Alpha* | *Delta* |
| Cat F | 12 | 490 | 257 | 601 | 5.90% | 0.90% | 4.10% |
|  |  | 146 | 229 | 303 |  |  |  |
| Cat G | 175 | 586 | 677 | 243 | 0.40% | 0.40% | 0.40% |
|  |  | 134 | 170 | 58 |  |  |  |
| Cat H | 94 | 687 | 825 | 2165 | 0.30% | 0.20% | 0.70% |
|  |  | 474 | 678 | 685 |  |  |  |
| Cat J | 175 | 627 | 719 | 247 | 0.30% | 0.40% | 0.40% |
|  |  | 318 | 241 | 79 |  |  |  |
| Cat L | 23 | 109 | 102 | 468 | -7.20% | 1.40% | 1.60% |
|  |  | 289 | 70 | 301 |  |  |  |

## Discussion

This study has demonstrated that the seroprevalence of anti-SARS-CoV-2 antibodies in the UK domestic cat population has increased over time, consistent with results ascertained in a survey of cats and dogs recently conducted in Canada (*49*) and the very low seroprevalence observed in the first and second waves of the pandemic in both Thailand and the UK (*7, 50*). This may be explained by the persistence of the humoral response over time with a consequent accumulation in the number of seropositives in the population. While increased seroprevalence during the later months of the pandemic may mean the likelihood of human-to-cat transmission is greater for newer variants that are more readily transmitted between humans (*51, 52, 53*), this has not been experimentally investigated.

Many samples collected at later timepoints had their highest titer against the ancestral B.1 or Alpha variants, despite the dominant circulating lineage in humans being either Delta or Omicron at the time (Fig 6). This may indicate the cats were infected during either the first or second (Alpha) wave of the pandemic and were not re-exposed during the subsequent Delta (third) or Omicron (fourth) waves, which is logical given the generally low overall seroprevalence. A relationship is evident between the proportion of seropositives, with respect to dominant variant, detected at different timepoints and the waves of infection in the UK’s human population. The dominant strain detected in cats appears to trail the emergence of each VOC into the human population, indicating repeated cross-species jumps between humans and cats and implicating owner-to-pet transmission as the primary route of infection, consistent with other serosurveys (*11, 54, 55, 56*).

Longitudinal samples were available from five seropositive animals in the study. In four cases, neutralizing antibody titers waned over time, similar to findings in studies of both infected and vaccinated humans (*34, 57*). Although a definitive protective threshold antibody level for SARS-CoV-2 has not yet been established, waning neutralizing antibody levels in humans post-vaccination has been associated with re-infection and reduced protection against novel variants (*35, 58, 59*).

Increasingly, mucosal immunity and neutralizing IgA are believed to play important roles in the anti-SARS-CoV-2 response, due to the virus infecting hosts via the respiratory tract (*60, 61*). Further investigation into the feline mucosal immune response against SARS-CoV-2 may paint a clearer picture of the impact of waning serum neutralizing antibody titers on susceptibility.

In the absence of sequence data, the variant to which the animal was exposed can only be inferred from serology, however, in some cases, the titer against the dominant variant was many times greater than the next highest titer, providing a strong case for it being the infecting variant. Three distinct patterns of immunity emerged according to which variant was neutralized most effectively, similar to previous findings in humans (*62*). It is likely the breadth and potency of variant-specific neutralization is influenced primarily by both the antigenicity of the variant, and the viral load post-infection. The trends observed for cats thought to have been infected with the B.1 variant are similar to the patterns of neutralization reported previously in humans (*40, 41*). It was shown that humans vaccinated with a Wuhan-Hu-1-based vaccine develop lower neutralization titers against the Delta (*63*) and Omicron (*64*) variants than against B.1 or Alpha. The distinct genetic and structural differences in the spike protein of Delta and Omicron could account for these variations in neutralizing antibody titers (*65, 66*).

As all samples tested in this study were collected prior to March 2022, none showed Omicron dominant neutralization. This finding was as anticipated since only a small proportion of samples were collected after the emergence of Omicron in the UK.

Although seropositivity indicates a cat has previously been infected with SARS-CoV-2, it is possible that a higher proportion of cats could have been infected with the virus but never developed or no longer have detectable neutralizing antibodies. Some human studies have identified small groups displaying either very low-level antibody responses post-vaccination (*67*) or no detectable response at all – many of these cases are thought to be correlated with underlying conditions or autoimmunity (*35*).

A higher proportion of pedigree cats were seropositive compared to their non-pedigree counterparts - this finding approached statistical significance. Pedigree cats are more likely to be indoor-only (*68*) and may therefore experience more close contact with their owners, meaning they are more exposed to SARS-CoV-2 if their owners become infected.

It should be noted that the sample population examined in this study, while broadly representative of the UK’s feline population, was inherently biased towards clinically sick animals. As all samples tested were remnants from diagnostic submissions, the majority of the animals would either have been showing signs of disease, newly rescued or under observation at the time of sampling (Fig 8). This means certain breeds that might be more susceptible to disease could have been overrepresented. For example, pedigree cats constitute approximately 10% of the UK feline population (*69*) but made up 31% of the samples included in this study, perhaps reflecting a higher morbidity in pedigree cats or increased willingness of pedigree cat owners to spend money on diagnostic testing. It is possible that SARS-CoV-2 seroprevalence could be higher in the population sampled, since cats attending veterinary clinics might be more likely to have genetic factors, immunosuppression or comorbidities which affect susceptibility to infection.

Our results demonstrate the importance of widespread testing of cats, to detect SARS-CoV-2 exposure and better understand the morbidity and mortality associated with infection in cats. Testing oropharyngeal swabs for SARS-CoV-2 RNA by RT-qPCR provides an opportunity to monitor for feline-specific mutations and transmission events from infected cats, as well as allowing for comparison with serology to accurately identify the causative variant of infection. Both widespread serological and qPCR-based testing are vital to address the One Health aspect of SARS-CoV-2 infection (*70*). Without further research to determine the importance of cats as a possible SARS-CoV-2 reservoir, a vital piece of the jigsaw may be missing in the attempt to bring global infections under control.

## Acknowledgements

The authors thank Dawn Dunbar^1^, Leigh Marshall^1^ and Andrea Bowie^1^ for assisting with sample provision, and the BBSRC and Serth and Gates Charity for funding this research. ^1^ Veterinary Diagnostic Service, School of Biodiversity, One Health and Veterinary Medicine, University of Glasgow.

## Supplementary Data

**Figure 8:**
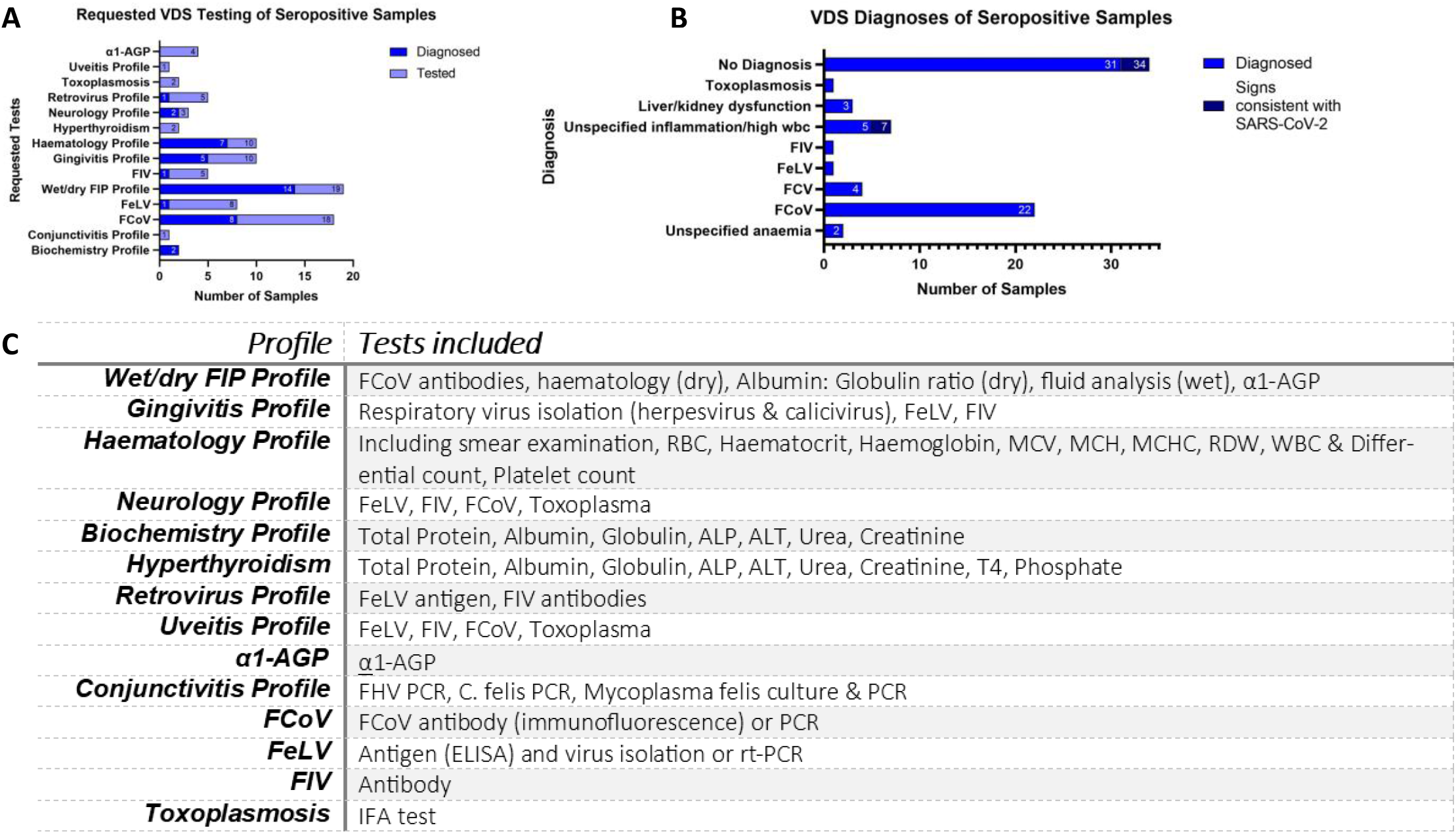
Overview of VDS tests conducted on dataset. All samples tested are residuals which had been submitted to the University of Glasgow’s Veterinary Diagnostic Service (VDS) for various tests. The tests requested for seropositive samples and whether these tests resulted in a diagnosis (A). The diagnoses of seropositive samples along with those exhibiting clinical signs consistent with human SARS-CoV-2 infections (Respiratory and GI symptoms and pyrexia) (B). The specific tests included in testing packages offered by the VDS (C).

